# Individual variability in structural brain development from late childhood to young adulthood

**DOI:** 10.1101/2021.02.04.429671

**Authors:** Kathryn L. Mills, Kimberly D. Siegmund, Christian K. Tamnes, Lia Ferschmann, Lara M. Wierenga, Marieke G. N. Bos, Beatriz Luna, Megan M. Herting

## Abstract

A fundamental task in neuroscience is to characterize the brain’s developmental course. While replicable group-level models of structural brain development from childhood to adulthood have recently been identified, we have yet to quantify and understand individual differences in structural brain development. The present study examined individual variability and sex differences in changes in brain structure, as assessed by anatomical MRI, across ages 8.0–26.0 years in 269 participants (149 females) with three time points of data (807 scans), drawn from three longitudinal datasets collected in the Netherlands, Norway, and USA. We further investigated the relationship between overall brain size and developmental changes, as well as how females and males differed in change variability across development. There was considerable individual variability in the magnitude of changes observed for all included brain measures. However, distinct developmental patterns of change were observed for total brain and cortical gray matter, cortical thickness, and white matter surface area, with individuals demonstrating either stability or decreases in early adolescence, then almost universal decreases during mid-to-late adolescence, before returning to more variable patterns in early adulthood. White matter volume demonstrated a similar developmental pattern of variability, but with individuals shifting from increases to a majority stabilizing during mid-to-late adolescence. We observed sex differences in these patterns, and also an association between an individual’s brain size and their overall rate of change. The present study provides new insight as to the amount of individual variance in *changes* in structural morphometrics from late childhood to early adulthood in order to obtain a more nuanced picture of brain development. The observed individual- and sex-differences in brain changes also highlight the importance of further studying individual variation in developmental patterns in healthy, at-risk, and clinical populations.

## Introduction

Longitudinal MRI research conducted over the past two decades has demonstrated that the human brain undergoes a prolonged course of development, with changes in morphometry observed in the cortex, as well as in white matter and subcortical structures, throughout childhood and adolescence (Aubert-Broche et al., 2013; Goddings et al., 2013; Lebel & Beaulieu, 2011; Mutlu et al., 2013; Raznahan et al., 2011; Reynolds et al., 2019; Vijayakumar et al., 2016; Wierenga, Langen, Oranje, et al., 2014; Wierenga, Langen, Ambrosino, et al., 2014; Wierenga, Bos, et al., 2018). While this research has had a profound impact on our understanding of brain development, most of these studies have focused on estimating group-level trajectories, and quantifying the degree of individual variability in structural brain development remains a neglected area of research (Becht & Mills, 2020). Characterizing variability across individuals in how the brain *changes* during development is needed to address some of the most pressing questions in developmental neuroscience. It is only with this knowledge that we can identify individuals who begin to deviate in neurotypical development, or tailor prevention and intervention efforts to impact the processes that are changing the most during different developmental periods.

A goal of developmental neuroscience is to define patterns of brain maturation. We know that, for instance, the cerebral cortex decreases in volume and thickness across the second decade of life before beginning to stabilize in the early twenties, and that cerebral white matter increases until some point in mid-to-late adolescence (Aubert-Broche et al., 2013; Mills et al., 2016; Tamnes et al., 2017; Wierenga, Langen, Oranje, et al., 2014). But the age at which these structures begin to stabilize—a measure of maturity for structural brain measures—likely varies across individuals. Subcortical structures also show heterogeneity in their volumetric development (Herting et al., 2018; Ostby et al., 2009; Wierenga et al., 2014), and our previous work has demonstrated that the amygdala and nucleus accumbens vary in when they reach a point of maturity across individuals (Mills et al., 2014). Identifying the periods of development when there is more individual variability in change vs. stability can inform theories on how immaturity vs. maturity in a given brain measure reflects cognitive, emotional, and behavioral processes across development (e.g. the imbalance model; Casey et al., 2008).

Several neurodevelopmental models of psychopathology hypothesize deviations in the rate of brain development for individuals at risk for, or with already developed, mental health disorders (Shaw et al., 2010). Neuroimaging studies have indeed demonstrated that rates of change in structural brain development relate to psychopathological symptoms (Bos, Peters, et al., 2018; Bos, Wierenga, et al., 2018; Ducharme et al., 2014; Muetzel et al., 2018; Whittle et al., 2020). Notably, sex differences are also present in patterns of structural brain changes across childhood and adolescence (Herting et al., 2014, 2018; Wierenga, Bos, et al., 2018; Wierenga, Sexton, et al., 2018), which may be pertinent to understanding sex differences in onset, prevalence, and progression of psychopathology (Paus et al., 2008; Shaw et al., 2010). Currently, there are several large-scale initiatives that aim to identify the genetic and environmental factors that shape the developmental course of the brain (e.g. ABCD, IMAGEN, Generation R). Moving forward, however, these investigations of brain development would benefit from knowing the periods of development when the most individual variability is likely to occur, and which measures of the brain demonstrate the most individual variability.

There are now several longitudinal MRI datasets of structural brain development that can be used to examine variability in brain change over time (Vijayakumar et al., 2018). Indeed, individual variability in how the brain develops can only be examined with longitudinal datasets. While large cross-sectional samples are able to quantify the normative range of values of a given brain measurement at different periods of development, only longitudinal datasets can quantify the normative range of how brain measurements change within individuals at different periods of development, as it is not possible to estimate average rates of change in a given developmental period with only one measurement collected per individual. Recently, we have conducted secondary analyses of multiple longitudinal datasets with two or more time points of data per individual to establish replicable group-level models of typical structural brain development from childhood to early adulthood (Herting et al., 2018; Mills et al., 2016; Tamnes et al., 2017). The focus of the present investigation is to characterize individual variability and sex differences in *changes* of brain structure (morphometry) across development. We do so by taking advantage of three separate datasets of developing individuals with three time points of data, which is necessary to model individual-level slopes in a multi-level analysis framework.

While the main focus of the present study is characterizing individual variability in structural brain changes, we also examine whether individuals who have higher or lower measurements of a given structure compared to similar-aged peers show different maturation patterns. This secondary focus of the present investigation is motivated by a common inference in the neuroimaging literature that the maturity of a developing individual’s brain size can be assessed by comparing them cross-sectionally to similar-aged peers. For example, children and adolescents with thinner cortex than similar-aged peers are often inferred as more mature, or faster-maturing, in their brain development (e.g., Paulus et al., 2019; Tamnes et al., 2018; Thijssen et al., 2020). In this example, the inference stems from the group-level observation that the cortex of the human brain decreases in thickness across childhood and adolescence, but fails to consider the large amount of variability neurotypically developing children can have in one given brain measure (Wierenga et al., 2019). For example, one child can have an average cortical thickness that is 85% of the value of another child the same age (from data presented in Tamnes et al., 2017). If the claim that children and adolescents with thinner cortex are more mature, we would expect that they would show less overall change in cortical thickness, as a lack of change in brain structure is one way to assess the maturity of the brain. We test this hypothesis, as well as the relationship between overall values of other structural brain measurements in an individual with observed changes during development. We also examine how the relationship between overall brain size and maturation patterns differ between females and males.

## Methods

### Participants

This study examined participants from three separate longitudinal datasets collected from independent research sites located in three countries: Leiden University (BrainTime), University of Oslo (Neurocognitive Development; NCD), and University of Pittsburgh (LunaCog). Only participants with three high-quality anatomical brain scans were included in the present analysis, for a total of 269 participants (149 females, 120 males). Demographic characteristics for each sample are described in Table 1, and the sampling design is illustrated in Figure 1. The distribution of ages varied slightly by dataset, with the average age of participants at first and last visit (and the interval between them) as follows: approximately 15–19 years (4 years) for BrainTime, 14–21 years (7 years) for NCD, and 15–18 years (3 years) for LunaCog. The number of data points included from females and males for each age (rounded to year) is illustrated in SFigure 1. Details regarding participant recruitment at each site are described in the Supplemental Material.

**Figure 1.**
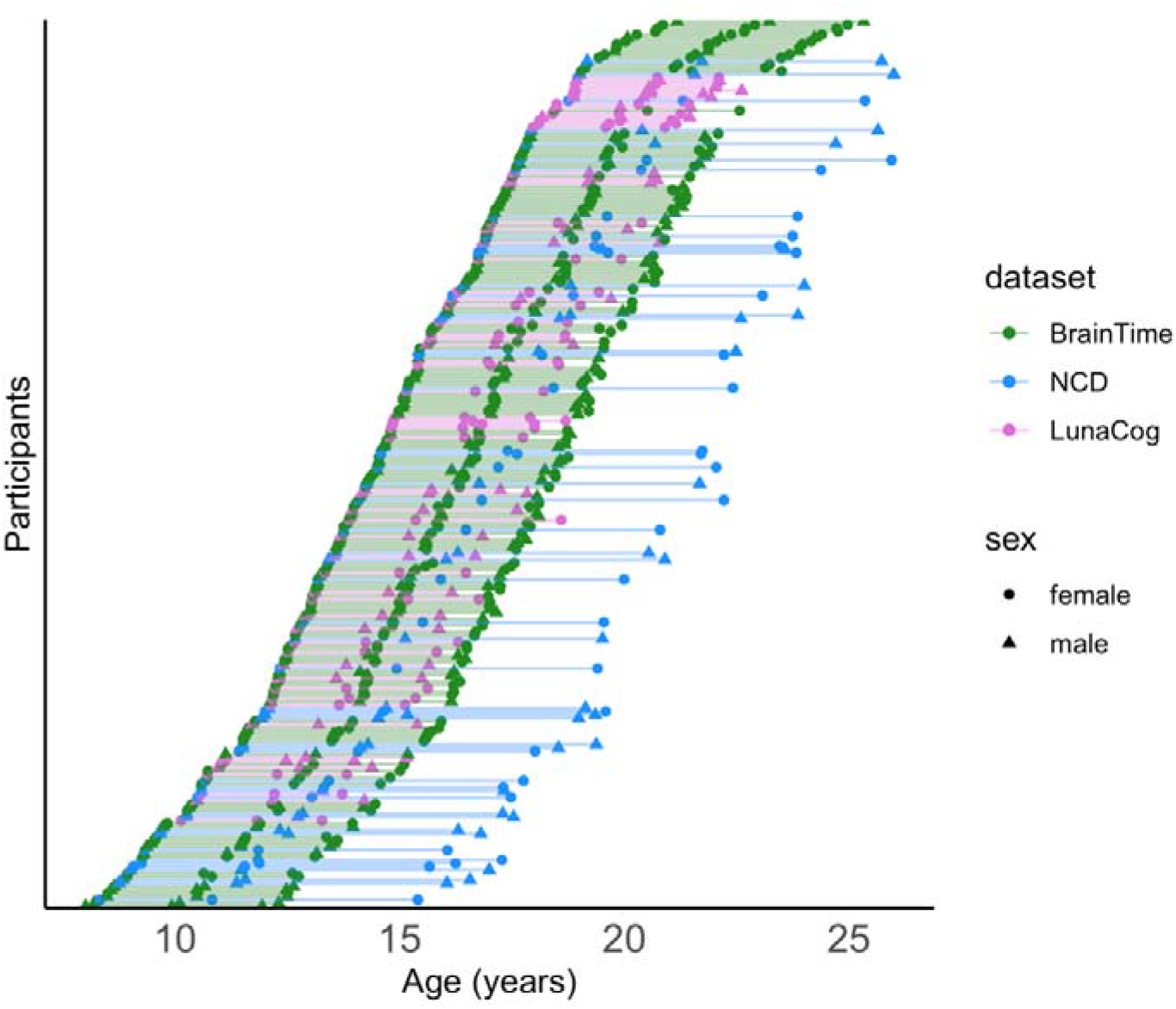
Scatter plot of age at scan for all participants. Each of the participants are shown in a different row, with each line connecting their three respective scans. Female (circle) and males (triangles) are denoted for each dataset (BrainTime: green; NCD: blue; LunaCog: Purple).

**Table 1.**
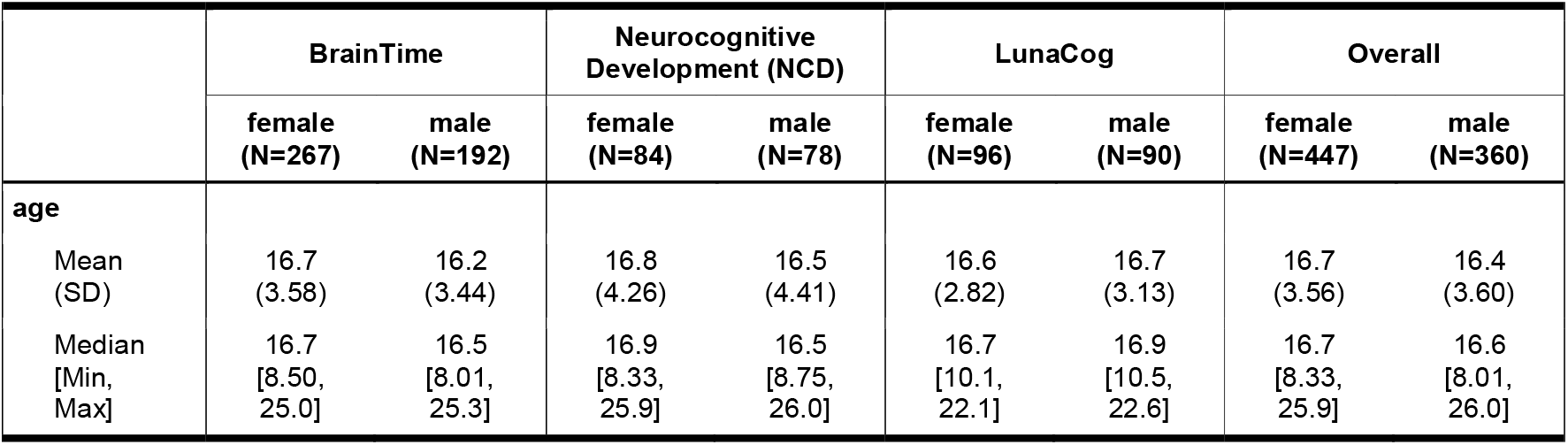
Distribution of age and sex for the number of scans for each dataset. All participants had to have three good-quality MRI scans to be included in the present study.

### Image processing

Participants in the BrainTime and LunaCog samples were scanned using 3-T MRI machines, while the NCD sample was scanned using a 1.5-T MRI machine. Details regarding image acquisition at each site are described in the Supplemental Material. MRI processing was performed with the FreeSurfer 6.0 image analysis suite, which is documented and freely available online (http://surfer.nmr.mgh.harvard.edu/), on workstations and operating systems at their respective universities (see Supplemental Material). The technical details of these procedures are described in detail in seminal publications (Dale et al., 1999; Fischl et al., 1999, 2002). This processing stream includes motion correction (Reuter et al., 2010), removal of non-brain tissue using a hybrid watershed/surface deformation procedure (Ségonne et al., 2004), automated Talairach transformation, non-parametric non-uniform intensity normalization (Sled et al., 1998), tessellation of the gray/white matter boundary, automated topology correction (Fischl et al., 2001; Ségonne et al., 2007), and surface deformation following intensity gradients to optimally place the gray/white and gray/cerebrospinal fluid borders at the location where the greatest shift in intensity defines the transition to the other tissue class (Dale et al., 1999; Dale & Sereno, 1993; Fischl & Dale, 2000). Each cortical model was registered to a spherical atlas using individual cortical folding patterns to match cortical geometry across participants (Dale et al., 1999).

Images were then processed using FreeSurfer 6.0’s longitudinal stream (Reuter et al., 2012). This process includes the creation of an unbiased within-participant template space and image using robust, inverse consistent registration (Reuter et al., 2010). Several processing steps, such as skull stripping, Talairach transforms, atlas registration as well as spherical surface maps and parcellations are then initialized with common information from the within-participant template, significantly increasing reliability and statistical power (Reuter et al., 2012). All images were assessed for quality, as described further in the Supplemental Material.

### Brain measures of interest

Measures of brain structure were computed at each time-point for each participant. For the purposes of this study, we included global measures of total gray matter volume, cortex volume, mean cortical thickness, white matter surface area, cerebral white matter volume, subcortical gray matter volume, as well as volumes of specific subcortical structures: amygdala, hippocampus, thalamus, pallidum, caudate, and putamen. We chose not to report on the nucleus accumbens of the FreeSurfer output, given less information about the test-retest reliability of the nucleus accumbens using the FreeSurfer longitudinal pipeline (Reuter et al., 2012).

### Analysis procedure

The first aim of the study was to characterize individual variability in structural brain change from late childhood into early adulthood. Each participant in the current analysis had three good-quality MRI scans, which allowed us to examine individual-level change across these three time points. We took two different approaches to measure variability in change in brain structure, which are described below.

First, we calculated change in a structure between each observation period, which gave us two observations of change per participant given that each participant had three time points of data. We then calculated the annualized change score by dividing the amount of change observed between time points by the amount of time between the two observation periods. We also calculated the annualized *percent* change, to provide an assessment of how a given structure changed relative to the overall size of the structure. We calculated the amount of change relative to the average size of the brain structure between observation periods. Specifically, this is how we calculated the annualized percent change for each structure for each of the two observation periods, with (*x* = observation):

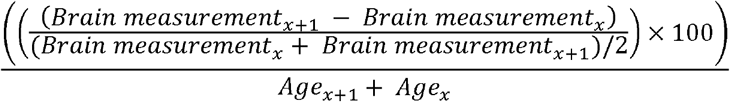

Our rationale for examining both annualized change and annualized percent change is so that we could assess if there were any notable differences between the two, as we have done in our previous work (Mills et al., 2016).

We categorized individuals as demonstrating an increase in a given brain measure if their annualized percent change was equal to or greater than the standard deviation of the annualized percent change observed in that particular measure across the sample, demonstrating a decrease if their annualized percent change was equal to or lesser than the negative value of the standard deviation, and stable if between these values. We categorized individuals with an equivalent method for the annualized change score.

For this first aim, we also applied a generalized additive mixture model (GAMM) to visualize the group-level developmental pattern of change across the age period, as well as to assess if these patterns differed between females and males. To assess sex differences, we compared three models: an age only model, a model including a main effect of sex and age, and a model including an interaction between sex and age. We compared these three models using Akaike Information Criterion (AIC) and likelihood ratio statistics (LR test) to avoid overfitting, selecting the model with the lowest AIC score that was significantly different from the more parsimonious models.

The second aim of the study was to examine the relationship between the rate of change of a given structural brain measure to an individual’s average structural brain measure of interest. Here, we took a different approach to measure variability in change in brain structure to better fit this aim. We first applied GAMM to model brain development as a function of age, with a participant-specific random intercept and slope. This is the same as fitting a hierarchical model for a given brain measurement (level 1) and slope (level 2). In level 1, the brain measure is modeled as a natural cubic spline with an intercept that varies by sex and dataset. In level 2, the slopes vary by sex and dataset and the deviation of participant’s mean outcome from the population mean. These can be fit into a single stage model by using interaction terms as follows:

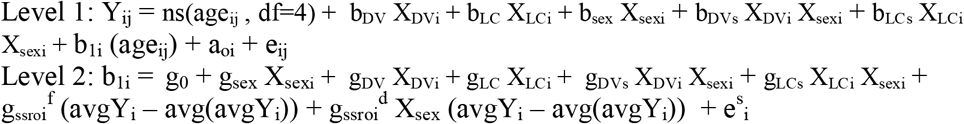

Where Y_ij_ is the brain outcome measure at the j^th^ visit (i.e. 1-3) for the i^th^ participant; X_DVi_, X_LCi_, and Xsex_i_ indicator variables for dataset (NCD, LunaCog with reference of BrainTime) and male sex, respectively; a_oi_ reflects the participant’s random intercept and e^s^_i_ their random slope; g_0_ is the slope of age in females from BrainTime (reference group), whereas g_sex_ reflects the difference in age slope in males versus females (also in BrainTime); g_DV_ and g_LC_ reflect the deviation in slopes in females between datasets (NCD, LunaCog with reference BrainTime); g_DVs_ and g_LSs_ capture the additional deviation in age slope in males vs females by dataset; and g_ssroi_^f^ and g_ssroi_^d^ are the primary covariates of interest; plus error (e_ij_). Specifically, gssroi captures the association between the participant’s slope and the difference in that slope as a function of the participant’s average brain outcome (avgY_i_) from the population average of the brain outcome of interest (avg(avgY_i_)) in females and g_ssroi_^d^ reflects how this differs in males and females. Each predictor variable is centered by subtracting the average value, with age_ij_ centered by participant. Variability in the outcome measure can introduce a negative bias into the association between rate of change and initial measure, also known as regression to the mean (Chiolero et al., 2013). To avoid this bias, we studied the association between rate of change and the participants’ average measure of the structure of interest. By comparing a participant’s slope with their average brain size across the three time points, we reduce the influence of measurement error in a participant’s intercept, as both the intercept and the slope will include the error from the baseline measurement, and this shared error could induce a spurious correlation when trying to study the relationship between the slope and intercept.

## Results

### Individual variability in structural brain change

We report the standard deviations for both the annualized change and annualized percent change across the sample for each structural brain measure in STable 1. Given that the annualized change and annualized percent change approaches resulted in almost identical developmental patterns of individual variability in structural brain change (compare Figure 2 with SFigure 2), we focus on describing the results from the annualized percent change measure since it allows for greater comparability across measures. For visualization, we plot annualized percent change against the age at the midpoint of observation period (Figure 2). To compare the number of individuals following these different patterns across late childhood to early adulthood, we binned individuals by yearly increments based on the individual’s age at the midpoint of observation period (Figure 3). Our GAM models examining annualized percent change by age are grouped together for visualization purposes in Figure 4.

**Figure 2.**
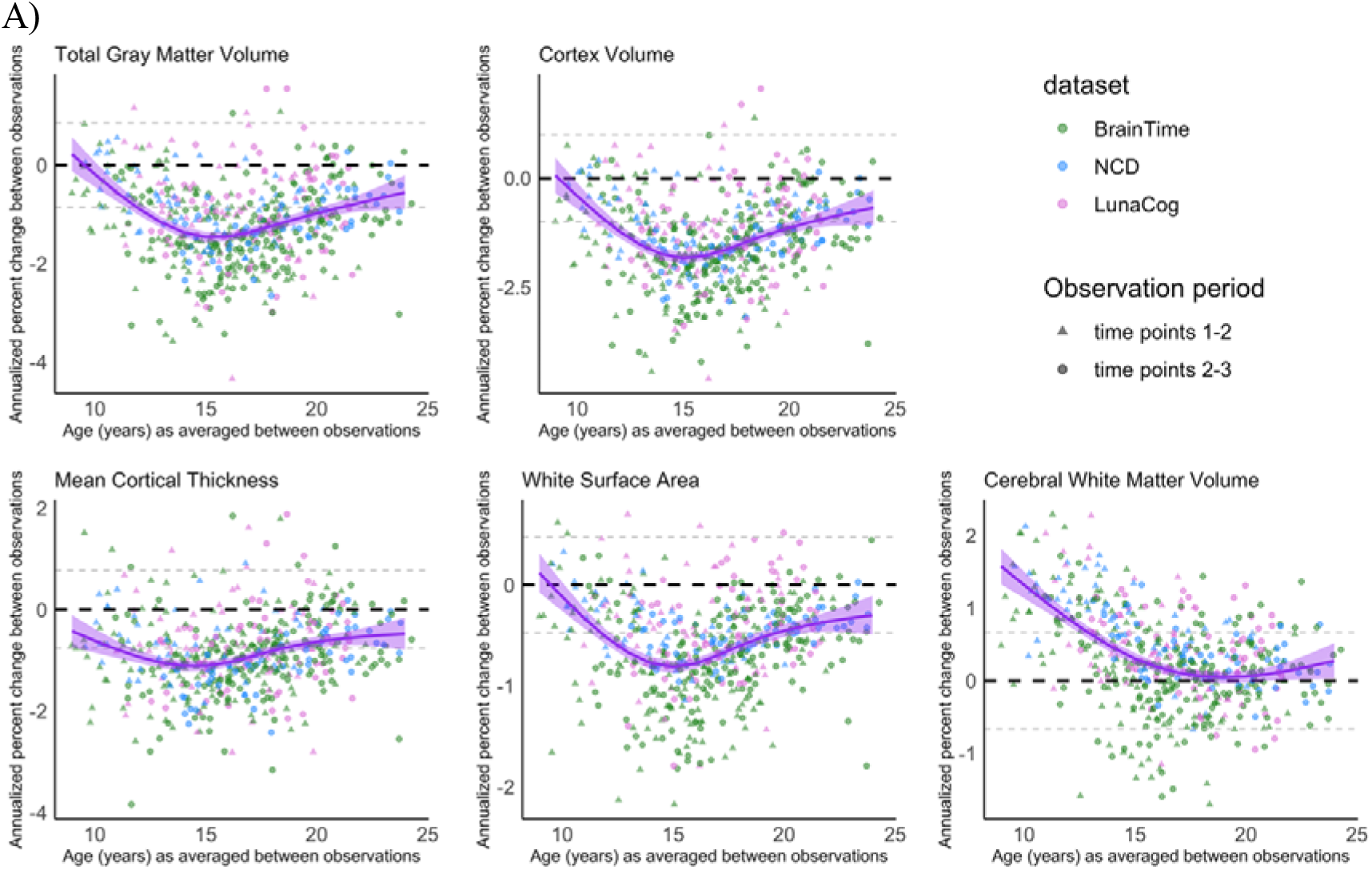
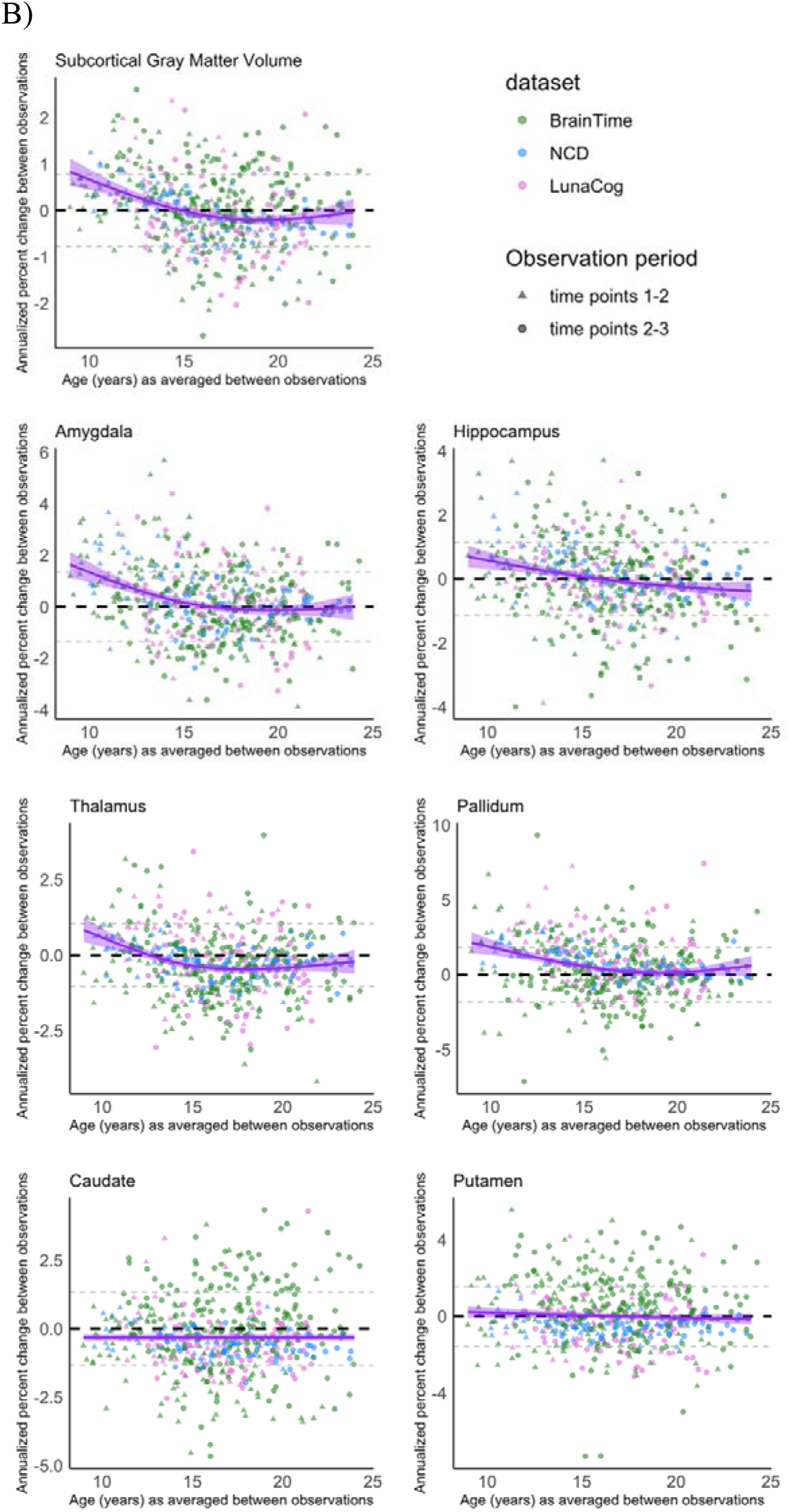
Variability in annualized percent change over time for each individual (y-axis), against age at the midpoint of the observation period. The purple line reflects group-level annualized percent change seen with age. The black dashed line marks 0 on the y-axis and the dashed gray lines represent the standard deviation of annualized percent change across the whole sample. A) Total gray matter volume, cortex volume, cortical thickness, white surface area, and white matter volume B) Subcortical gray matter volume and specific subcortical structures. The equivalent graphs for annualized change can be seen in SFigure2.

**Figure 3.**
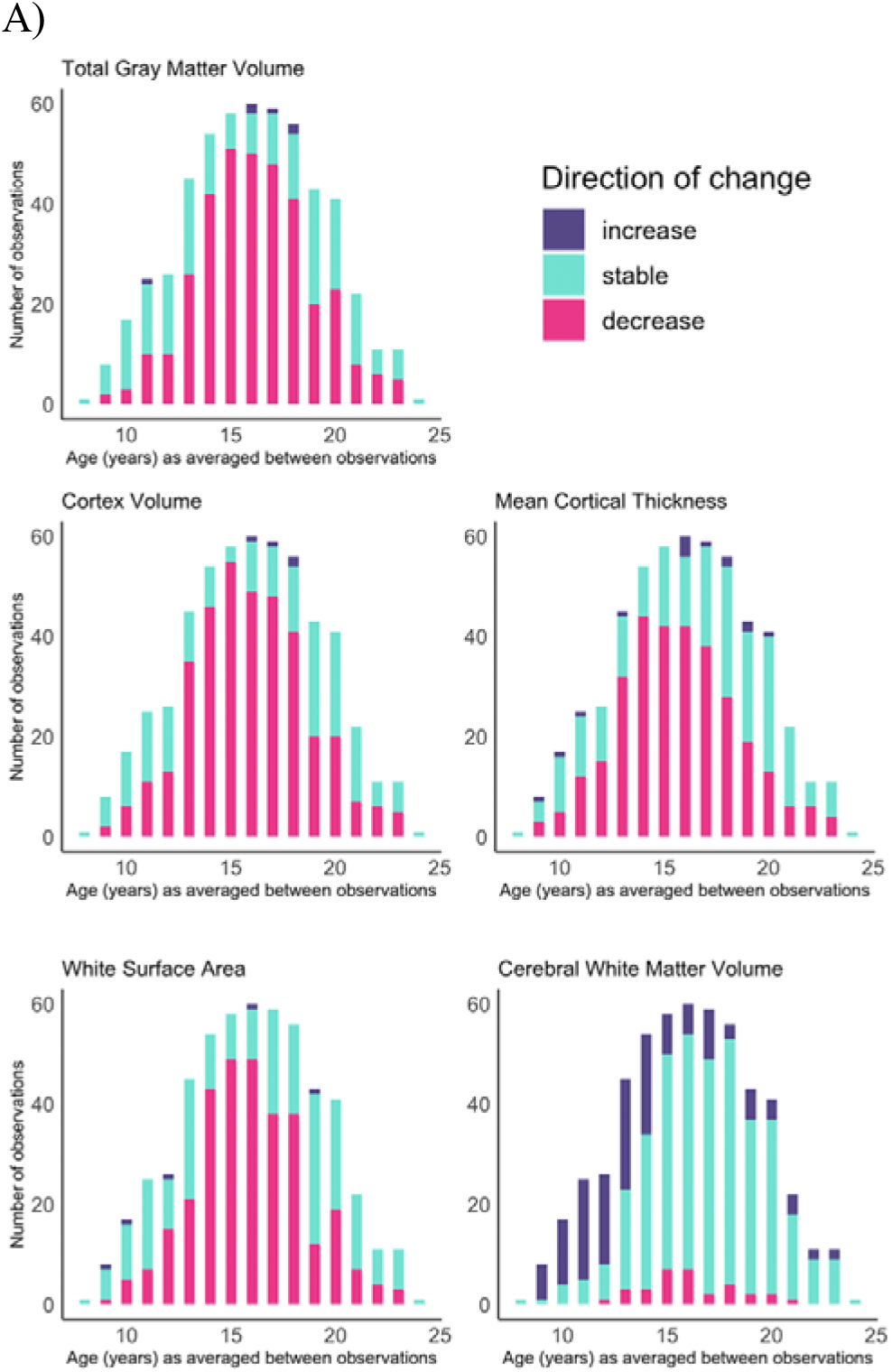
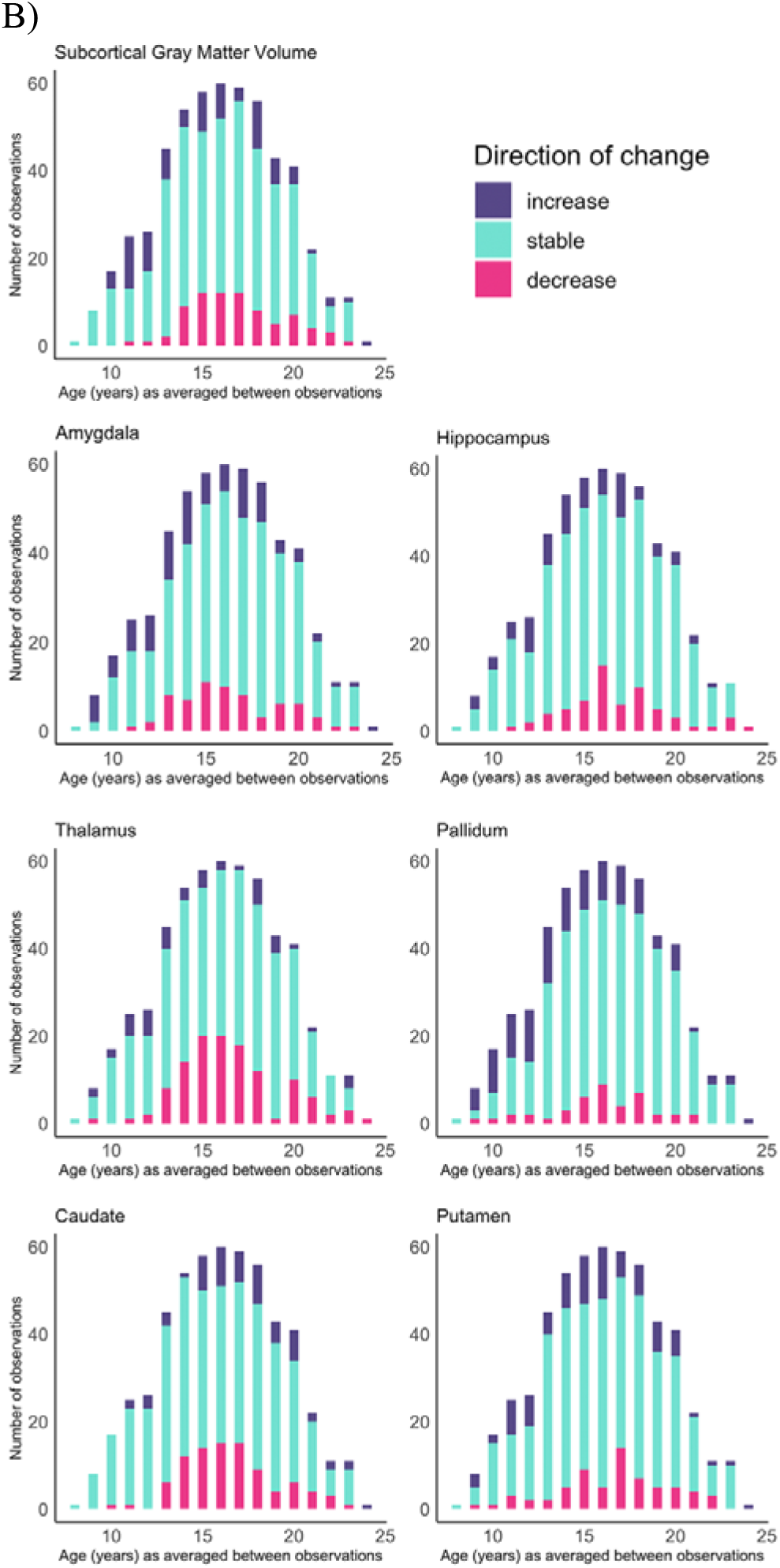
Number of individuals showing volume increases (purple), decreases (pink), or no change (turquoise) based on the individual’s age at the midpoint of observation period. A) Total gray matter volume, cortex volume, cortical thickness, white surface area, and white matter volume B) Subcortical gray matter volume and specific subcortical structures. See SFigure 3 for the same data graphed by percentage of individuals in each category.

**Figure 4.**
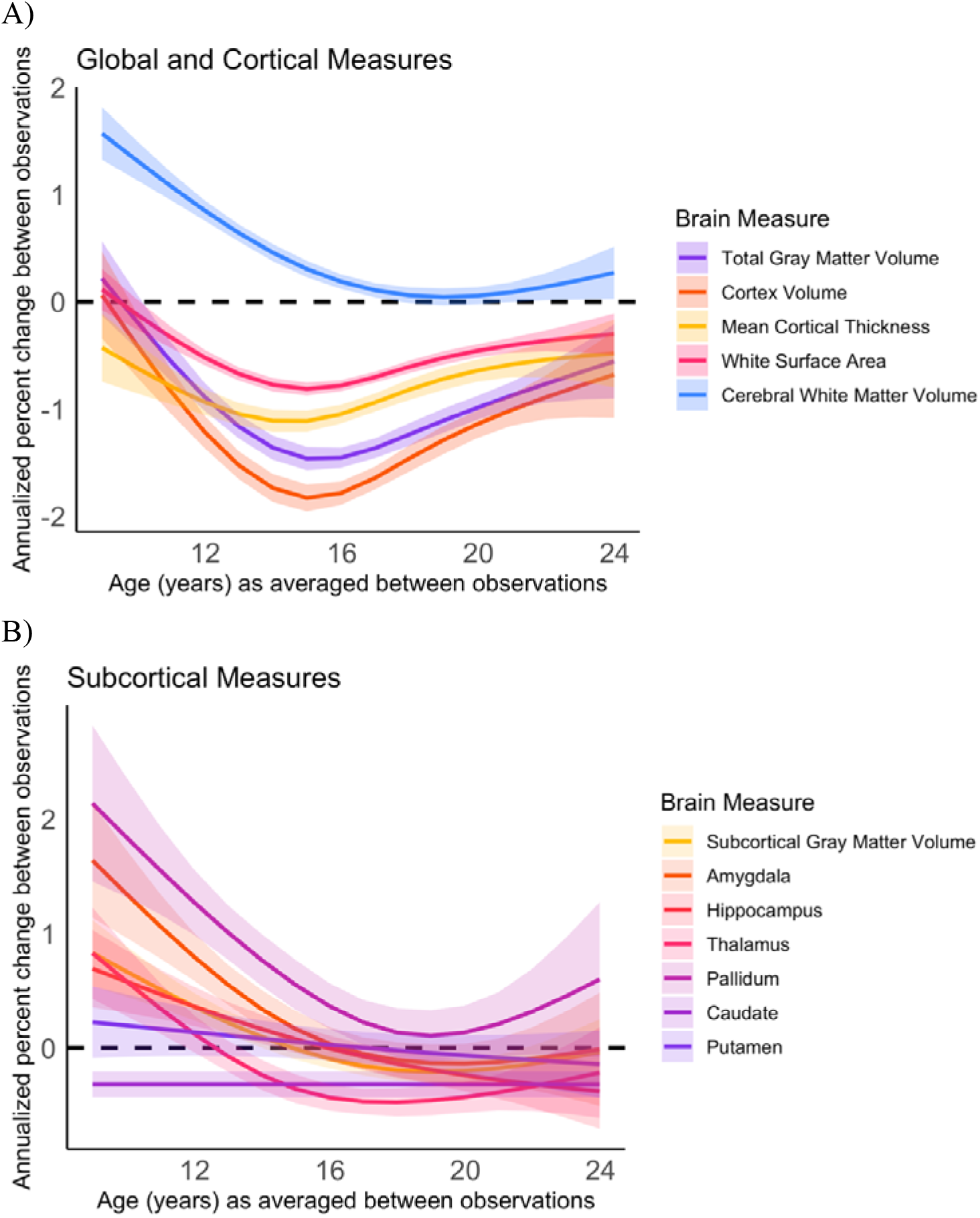
GAM models for annualized percent change for multiple brain measures together on one graph for the purpose of comparing group-level patterns of change. A) Total gray matter volume, cortex volume, cortical thickness, white surface area, and white matter volume B) Subcortical gray matter volume and specific subcortical structures.

Individual variability in direction and magnitude of change was present to some degree for every structural brain measure examined, and this variability was present across late childhood and into early adulthood. However, distinct developmental patterns of change were observed for total brain and cortical gray matter, mean cortical thickness, and white matter surface area. The majority of individuals demonstrated either stability or decreases in early adolescence for cortical measurements and total gray matter volume, whereas in mid-adolescence (roughly between ages 14-17 years), the majority of individuals showed decreases in these measurements. By late adolescence and into early adulthood, there was again more variability, with the majority of individuals demonstrating either stability or decreases. These cortical measurements also demonstrated similar patterns of change, with the largest magnitude in decrease in change observed for most individuals in the early to mid-teens, and more stability observed in the early twenties (Figure 4). For cerebral white matter volume, the majority of individuals demonstrated increases in early adolescence, shifting to the majority demonstrating stability by mid-to-late adolescence. For subcortical gray matter volume and specific subcortical structures, variability in direction and magnitude of change was visible throughout the age-range investigated. For some structures, there appears to be subtle developmental patterns for directions of change, e.g. for the pallidum, most individuals demonstrated either an increase or stability in early adolescence and no substantial change from mid-adolescence onward.

We observed sex differences in the overall magnitude of annualized percent change for total gray matter volume, white surface area, cerebral white matter volume, subcortical gray matter volume, and for the pallidum, and we observed sex differences in the magnitude and pattern of annualized percent change for cortex volume (Figure 5). The selected best fitting model did differ for two brain measures when change was assessed as annualized change instead of annualized percent change (see STable 3). For total gray matter volume, the model including sex and age as an interaction was the best fitting model when annual percent change was modeled by age, whereas this interaction model did not converge when annualized percent change was modeled by age. Further, the main effect of sex observed for white surface area when annualized percent change was modeled by age was not present when annual percent change was modeled by age. Model comparison statistics for both annualized change and annualized percent change measures are detailed in STable 3.

**Figure 5.**
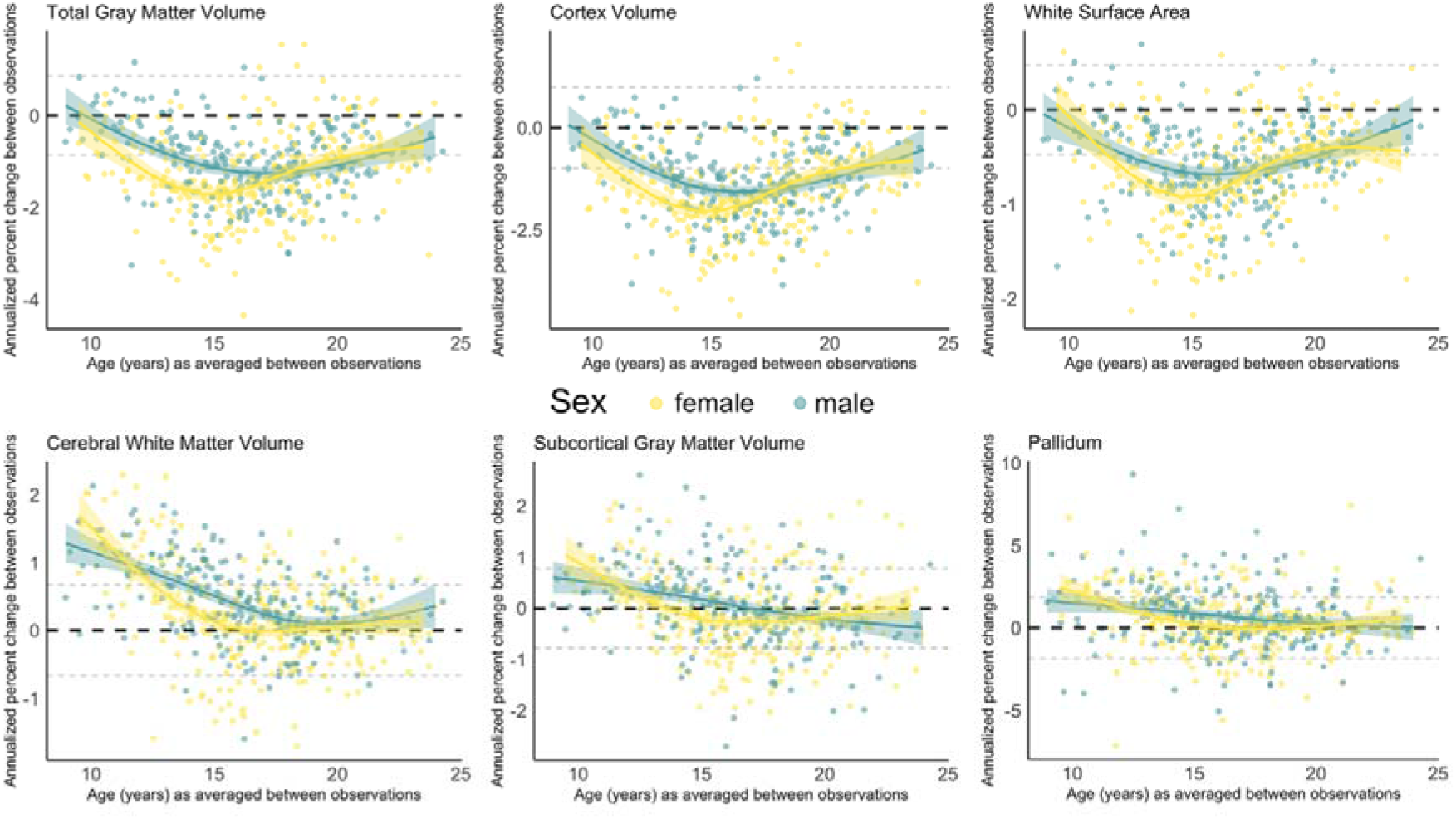
GAM models for annualized percent change for the brain measures where a model including sex either as a main or interactive effect was selected as a better fitting model based on likelihood ratio tests and AIC (see STable 3 for model comparison statistics). For total gray matter volume, white surface area, cerebral white matter volume, subcortical gray matter volume and pallidum, the best fit model included sex as a main effect. For cortex volume, the best fitting model included an interaction between sex and age. Female participants and their best fitting GAM model are represented in yellow, whereas males are represented in green.

### Individual differences in brain size and rates of change

Results for the relationship between the rate of change of a given structural brain measure to an individual’s average brain measurement and their sex are summarized in Table 2. Complete model outputs for each cortical and subcortical outcome can be found in STable 4. Individual’s rates of change were found to relate to an individual’s total gray matter volume, cortex volume, mean cortical thickness, and white matter surface area differently between females and males (Figure 5). Females with larger brain measurements showed greater negative rates of change (i.e. steeper decreases) for total gray matter and cortex volumes, mean cortical thickness, white surface area as compared to females with smaller brain measurements. However, males showed similar rates of change regardless of brain measurement size. Individual differences in rate of change were not related with an individual’s average brain size for cerebral white matter volume, subcortical gray matter volume, or any of the specific subcortical structures examined in the present study (Table 2).

**Table 2.** Generalized Additive Mixed-effects Models (GAMM) examining associations between participant’s slope and average brain size. Degrees of freedom, F-statistic, and p-value for Type III sum of squares are presented for each fixed effect of interest, as well as total observations and overall model fit (R^2^).

| *Measure* | *Finding* | *Individual Slope by Brain Size* |  |  |  |  |  |  |
| --- | --- | --- | --- | --- | --- | --- | --- | --- |
| | | *Effect*<br>*(in Females)* | | | *Sex Difference*<br>*(vs. Male)* | | | $R^2$ |
|  |  | *df* | *F* | *p* | *df* | *F* | *p* |  |
| Total Gray Matter Volume | Sex Interaction | 1 | 26.488 | **0.000000406** | 1 | 12.801 | **0.000386** | 0.988 |
| Cortex Volume | Sex Interaction | 1 | 28.635 | **0.000000143** | 1 | 9.764 | **0.0019** | 0.987 |
| Mean Cortical Thickness | Sex Interaction | 1 | 4.105 | **0.04331** | 1 | 6.734 | **0.00975** | 0.924 |
| White Surface Area | Sex Interaction | 1 | 9.905 | **0.0018** | 1 | 3.846 | **0.051** | 0.997 |
| Cerebral White Matter Volume | None | 1 | 1.062 | 0.303 | 1 | 0.576 | 0.448 | 0.996 |
| Subcortical Gray Matter Volume | None | 1 | 2.542 | 0.111436 | 1 | 2.626 | 0.105767 | 0.982 |
| Amygdala | None | 1 | 0.233 | 0.630 | 1 | 1.284 | 0.258 | 0.975 |
| Caudate | None | 1 | 0.508 | 0.476 | 1 | 0.123 | 0.726 | 0.972 |
| Hippocampus | None | 1 | 1.36 | 0.244 | 1 | 1.36 | 0.244 | 0.980 |
| Pallidum | None | 1 | 2.086 | 0.149 | 1 | 1.608 | 0.205 | 0.949 |
| Putamen | None | 1 | 0.006 | 0.940 | 1 | 0.047 | 0.828 | 0.956 |
| Thalamus | None | 1 | 0.002 | 0.965 | 1 | 0.017 | 0.897 | 0.980 |

## Discussion

The current collaborative research study utilized three longitudinal datasets including a total of 807 scans from 269 participants to quantify individual variability in the development of whole brain, cortical, and subcortical measurements across 8.0 to 26.0 years of age. Individual variability was present in each examined brain measurement, demonstrating that even when the majority of individuals follow a certain development pattern, some individuals will differ in the *direction* and *magnitude* of change. The majority of individuals demonstrated decreases in total gray matter volume, cortex volume, mean cortical thickness, and white matter surface area in mid-adolescence, with more variability present during the transition into adolescence and the transition into early adulthood. In contrast, the majority of individuals demonstrated increases in cerebral white matter in the transition into adolescence, with the majority showing stability starting in mid-adolescence and continuing into adulthood. Most individuals demonstrated little to no substantial change in subcortical structures or overall subcortical gray matter volume across the age range examined. Sex differences were largely reserved to cortical and whole brain measurements, in both our examination of patterns of annualized percent change across age, as well as in the association between brain structure size and rate of change.

Given that the majority of longitudinal development MRI studies in children and adolescents have been limited by two time points of data, most studies have looked at group-level effects of age. Few studies, however, have examined individual differences in how these structures change over time. The present study adds to the existing research by details just how variable changes in brain structure can be across late childhood and into young adulthood. Not only do individuals vary substantially in their overall brain structure size, but they can also differ in the magnitude, and sometimes direction, of observed developmental changes. We identified points in development where there is more individual variability in direction of change for specific structural measurements, which can inform future work in a number of ways. For example, research that aims to examine linear developmental processes in cortical measurements may want to constrain their age range to the teen years as this is when we can have the highest confidence that most individuals will be showing the same direction of change. Furthermore, researchers might choose to examine cortical volume over cortical thickness or surface area for such an investigation, as this cortical measurement shows the most consistency in direction of change during the teen years. The current study can also inform research wishing to assess individual differences in “inflection points” or brain maturation. Notably, for cerebral white matter volume, the majority of variability was in regards to when an individual begins to show stability in that measurement.

Examining the relationship between an individual’s brain size and overall developmental change revealed differing patterns between females and males. For example, females with larger total gray matter and cortical volume, cortical thickness, and cortical surface area measurements also demonstrated larger decreases in those measurements across time. This association was not present in males. These findings provide additional evidence suggesting that not only do individual differences exist in the overall level and pattern of development (i.e. slope), but also that these two properties of individual differences in cortical brain development relate to one another in a region-specific fashion.

The observed individual-level and sex differences in brain *changes* in from late childhood into young adulthood highlight the importance of further studying individual-level trajectories of brain maturation in cognitive and clinical neurodevelopmental investigations. A closer examination of each participant’s change over time suggests a complex interplay as to where the individual may fall relative to the group-average brain measurement. Regardless, if trajectories of structural brain development are linked to overall size or not for any given brain region, the amount of individual variability seen among each of these structures demonstrates that prior work aimed at understanding risk for psychopathology between the sexes via group-level sex differences in rates of brain maturation may miss the mark in identifying which individuals go on to develop mental health problems.

### Limitations

A substantial limitation to the current investigation is the lack of sociodemographic information available across the three longitudinal samples included in our analysis, which limits the ability to explore how potential environmental factors contribute to the variation seen in structural brain development and how this, in turn, may relate to behavior. Further, our definition of the threshold for what constituted a developmental change in a given brain measure represents a best estimate rather than a definitive rule. We calculated this threshold for each brain measure based on the overall standard deviation in annualized change (or annualized percent change) observed across the entire sample, which yielded estimates similar to those calculated from test-retest reliability studies of FreeSurfer measurements (Morey et al., 2010; Reuter et al., 2012). Nevertheless, our threshold for identifying change could have an impact on the conclusions of the present study and we suggest that future studies include multiple scans per individual in a given time point in order to best differentiate between time point measurement error from true developmental change. Finally, future work examining individuals with at least four time points of data would allow for estimation of non-linear individual slopes, which is more likely to resemble the shape of development for many structural brain measurements spanning the period of late childhood to early adulthood.

### Conclusions

The present study demonstrates that individuals vary in the *direction* and *magnitude* of structural brain changes across late childhood into young adulthood. While, at the group-level, cortical volume, thickness, and surface area followed similar patterns of change, with the greatest magnitude of decreases observed in the early to mid-teens, similar-aged participants could demonstrate very different rates of change across the age span examined. The magnitude of changes observed differed between females and males for whole brain measurements. Individual variability in rates of change related to overall brain size differently between females and males. Female participants demonstrated a negative relationship between brain size and change for cortical measurements, whereas this pattern was not seen in males.

**Figure 6.**
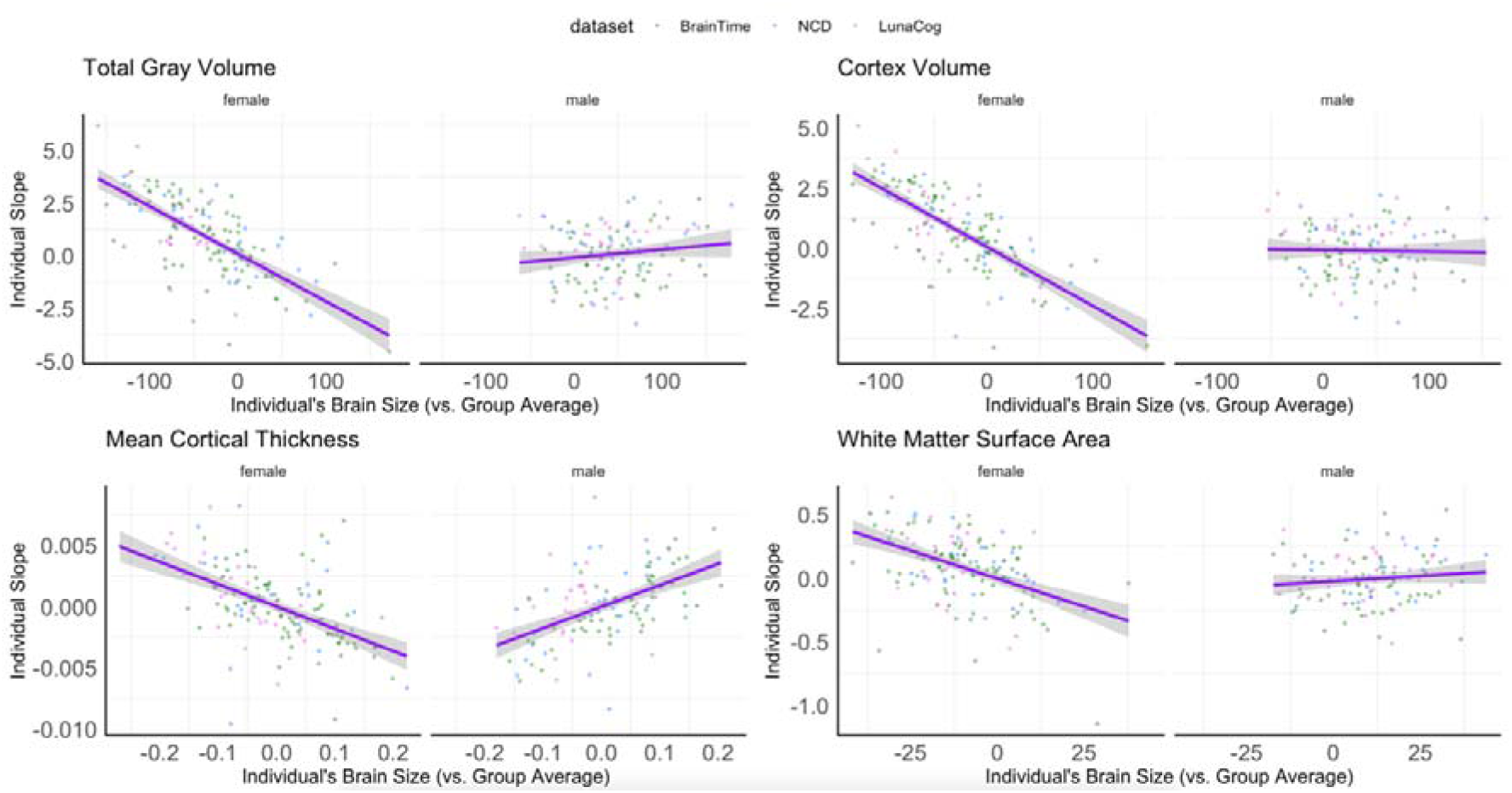
Significant sex differences in the association between individual’s slope as a function of an individual’s average brain measure compared to the group average (i.e. based on their study site and sex). Graphs present the magnitude of slope (i.e. rate of change) based on if an individual’s relative brain size to the group, with negative values reflecting smaller volumes and positive values reflecting larger volumes. Note, volume estimates have been scaled by 1000.

## Acknowledgements

K.L.M., C.K.T., L.F. and M.G.N.B were supported by the Research Council of Norway (RCN) grant number 288083. C.K.T was also supported by the RCN grant number 288083 and the South-Eastern Norway Regional Health Authority grant number 2019069. M.M.H was supported by the National Institutes of Health under award number K01 MH1087610. L.M.W was supported by the European Council starting grand scheme (ERC-2010-StG_263234 to E.A. Crone).

