## Supplemental Materials for "Individual variability in structural brain development from late childhood to young adulthood"

**S1. Methods**

Below we present sample descriptions for each sample. We do not include details about race and ethnicity in the samples because this information was not available for all samples. Further, due to differences in research ethics and standards, as well as in the goals and designs of the different studies, sociodemographic factors are not necessarily operationalized in the same manner across samples.

S1.1 Participants

S1.1.1 BrainTime sample

The BrainTime sample (Achterberg et al., 2016; Braams et al., 2015; Braams & Crone, 2016; Peters et al., 2016; van Duijvenvoorde et al., 2016) included 153 right-handed participants (89 females) between 8 and 23 years old, who were recruited in Leiden, the Netherlands through local schools and advertisements and had three high-quality scans. None of the participants reported a history of neurological or psychiatric disorders. All participants (or participant's primary caregivers for minors) provided written informed consent. Adults were paid for their participation, and minors and their primary caregivers received presents and a fixed payment for travel reimbursement. The internal review board from the Leiden University Medical Center approved the study procedures.

S1.1.2 Neurocognitive Development (NCD) sample

The Neurocognitive Development sample (Ferschmann et al., 2019; Tamnes et al., 2010, 2013) included 54 participants (28 female) who were recruited though newspaper advertisements and local schools in Oslo, Norway and had three high-quality scans. Written informed consent was obtained from all participants older than 12 years of age and from a parent of participants under 16 years of age, while participants under 12 years of age gave oral informed assent. At each time points, parents and participants aged 16 years or older complete screening for each participant with separate standardized health interviews to ascertain eligibility. Participants were required to be right-handed, fluent Norwegian speakers, have normal or corrected-to normal vision and hearing, not have a history of injury or disease known to affect central nervous system (CNS) function, including neurological or psychiatric illness or serious head trauma, not be under psychiatric treatment, not use psychoactive drugs known to affect CNS functioning, not have had complicated or premature birth, and not have MRI contraindications. The project was approved by the Regional Committee for Medical and Health Research Ethics.

S1.1.3 LunaCog sample

The LunaCog sample (Geier & Luna, 2012; Paulsen et al., 2015) sample included 62 participants (32 female) who were recruited from Pittsburgh and surrounding areas for behavioral testing and neuroimaging approximately every 15 months for two-and-a-half years and who had three high quality scans. Participants or their legal guardians provided informed consent or assent prior to participating in this study and all experimental procedures were approved by the Institutional Review Board at the University of Pittsburgh.

S1.2. Image Acquisition

S1.2.1 BrainTime Sample

All participants were scanned on a 3-Tesla whole body Philips Achieva MRI system (Best, The Netherlands). High-resolution 3D T1-weighted anatomical scan were obtained: TR 9.75 ms; TE 4.59 ms; FA 8º; FOV 224 x 168 x 177.333; 140 slices; voxel size .8756 × .875 × 1.2 mm. All anatomical scans were reviewed and cleared for gross abnormalities by a radiologist. The scans were processed on the same operating system (Linux Centos 6.3) and workstation.

S1.2.2 NCD Sample

All participants were scanned with a 12-channel head coil on the same 1.5-T Siemens Avanto scanner (Erlangen, Germany). The pulse sequence used for morphometric analyses was a 3D T1-weighted MPRAGE with the following parameters: TR 2,400 ms; TE 3.61 ms; inversion time (TI) 1,000 ms; FA 8°; matrix 192 × 192; field of view 240; 160 sagittal slices; voxel size 1.25 × 1.25 × 1.20 mm. The sequence was repeated at minimum twice in each session. Each scan took 7 min 42 s. A clinical neuroradiologist evaluated all scans for gross abnormalities.

S1.2.3 LunaCog Sample

All participants were scanned on a 3.0-T Siemens Allegra scanner at the Brain Imaging Research Center, University of Pittsburgh, Pittsburgh, PA. High-resolution anatomical data were collected using a magnetization prepared rapid acquisition gradient-echo (MP-RAGE) pulse sequence with 192 slices (1-mm slice thickness) in the sagittal plane. These data were processed for the current analysis on the University of Oregon high performance computer, Talapas. Each image was inspected for motion and reconstruction errors by a trained research assistant, and scans with motion or reconstruction errors were removed from the present analysis.

**S2. Results**

**STable 1** Standard deviations (SD) of the observed in that particular measure across the sample for each structure examined in the present investigation.

| **Structure Name** | Annualized percent change SD | Annualized change SD |
| --- | --- | --- |
| Total Gray Matter Volume | 0.86 | 6344.56 |
| Cortex Volume | 1.00 | 5563.69 |
| Mean Cortical Thickness | 0.77 | 0.02 |
| White Surface Area | 0.47 | 875.32 |
| Cerebral White Matter Volume | 0.67 | 3004.34 |
| Subcortical Gray Matter Volume | 0.77 | 515.56 |
| Amygdala | 1.35 | 47.25 |
| Hippocampus | 1.14 | 107.19 |
| Thalamus | 1.03 | 169.4 |
| Pallidum | 1.83 | 77.16 |
| Caudate | 1.33 | 115.21 |
| Putamen | 1.57 | 191.6 |

**STable 3a** Model comparisons for GAM Models of annualized absolute and annualized percent change for total gray matter volume, cortex volume, cortical thickness, white surface area, and cerebral white matter volume. We compared three models: an age only model (age), a model including age and a main effect of sex (main) and a model including main effects of age and sex as well as an interaction between age and sex (interaction). The best fitting model as determined by our model comparison criteria (lower AIC and significantly different at alpha 0.05 from less complex models) are highlighted in yellow. AIC= Akaike Information Criterion. BIC= Bayesian Information Criterion.

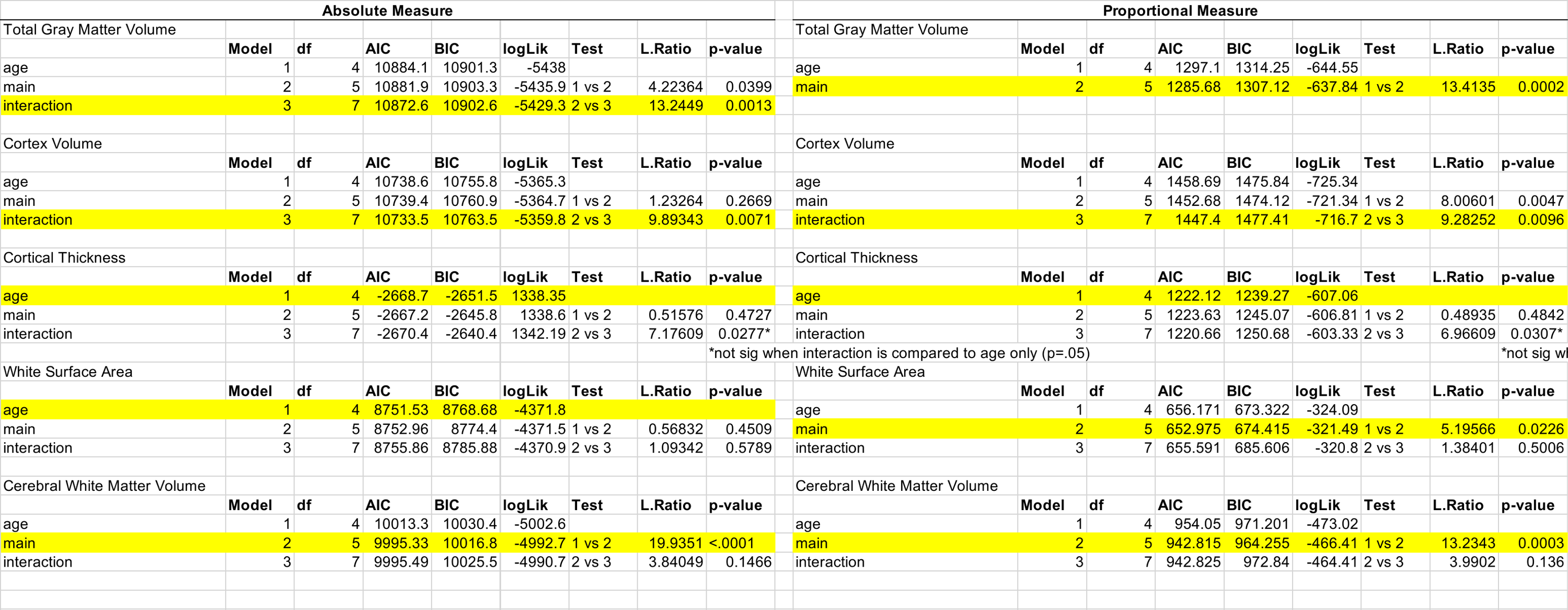

**STable 3b** Model comparisons for GAM Models of annualized absolute and annualized percent change for subcortical gray matter volume and subcortical structures. We compared three models: an age only model (age), a model including age and a main effect of sex (main) and a model including main effects of age and sex as well as an interaction between age and sex (interaction). The best fitting model as determined by our model comparison criteria (lower AIC and significantly different at alpha 0.05 from less complex models) are highlighted in yellow. AIC= Akaike Information Criterion. BIC= Bayesian Information Criterion.

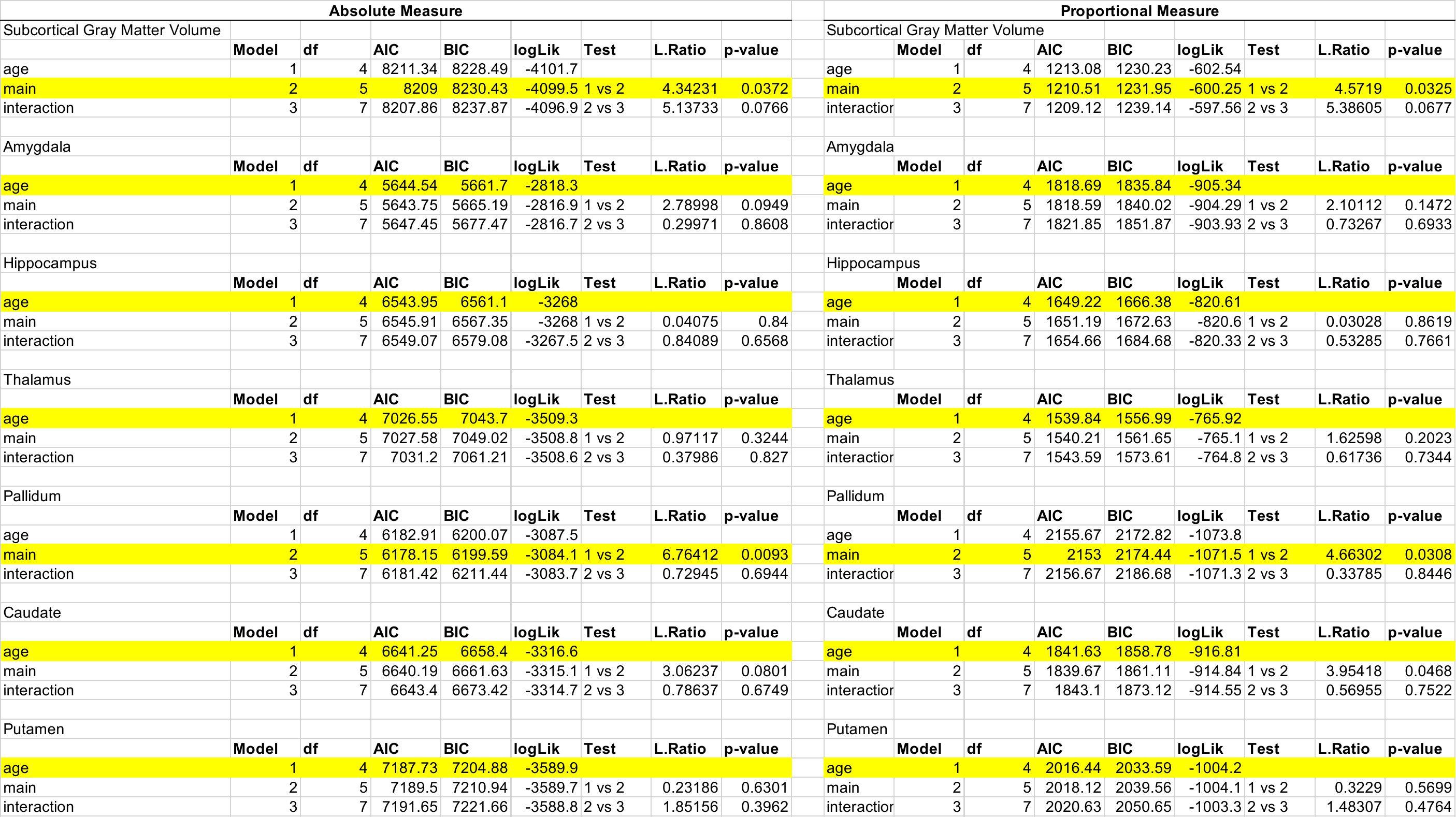

**STable 4a** GAM model estimates for total cortical, subcortical, and whole brain metrics for analysis comparing overall brain size to change. Beta coefficient estimates, confidence intervals, and p-values provided by fixed effects and edf and p-value for random effects term of participant’s intercept and slope.

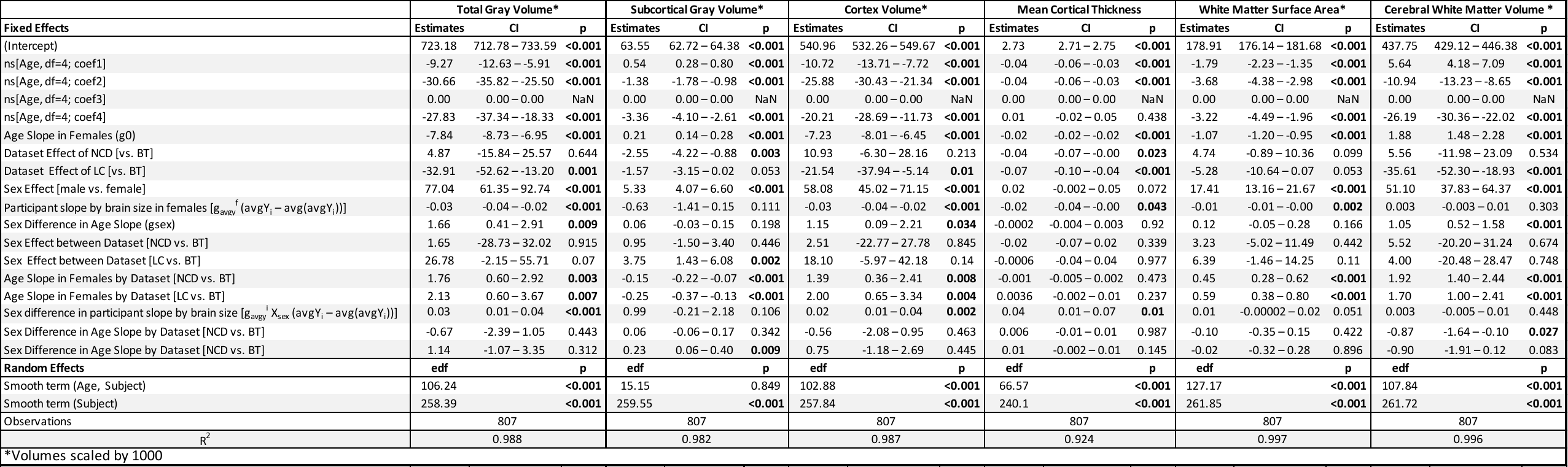

**STable 4b** GAM model estimates for each predictor in models for subcortical regions of interest for analysis comparing overall brain size to change. Beta coefficient estimates, confidence intervals, and p-values provided by fixed effects and edf and p-value for random effects term of participant’s intercept and slope.

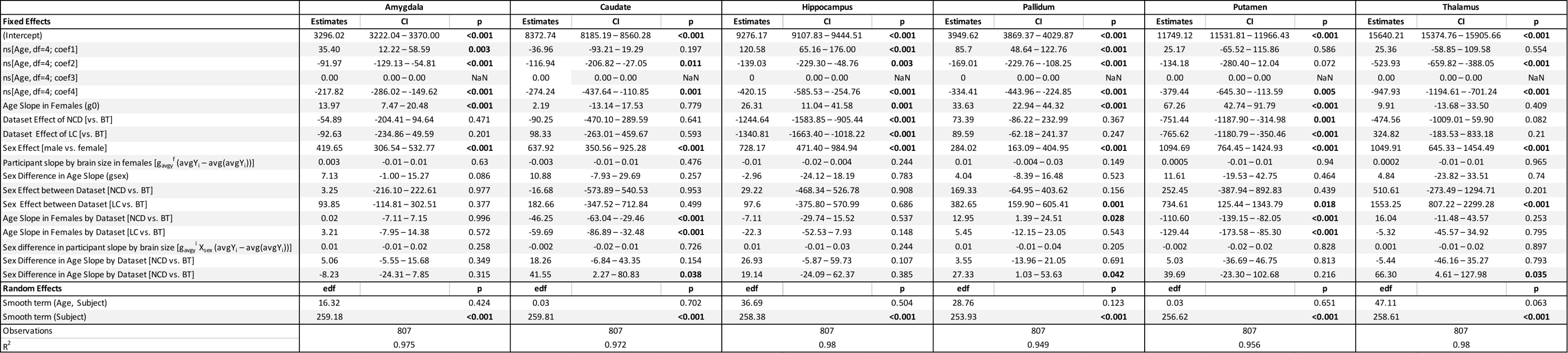

**SFigure 1**. Histogram illustrating the number of scans from female and male participants per age, binned by years.

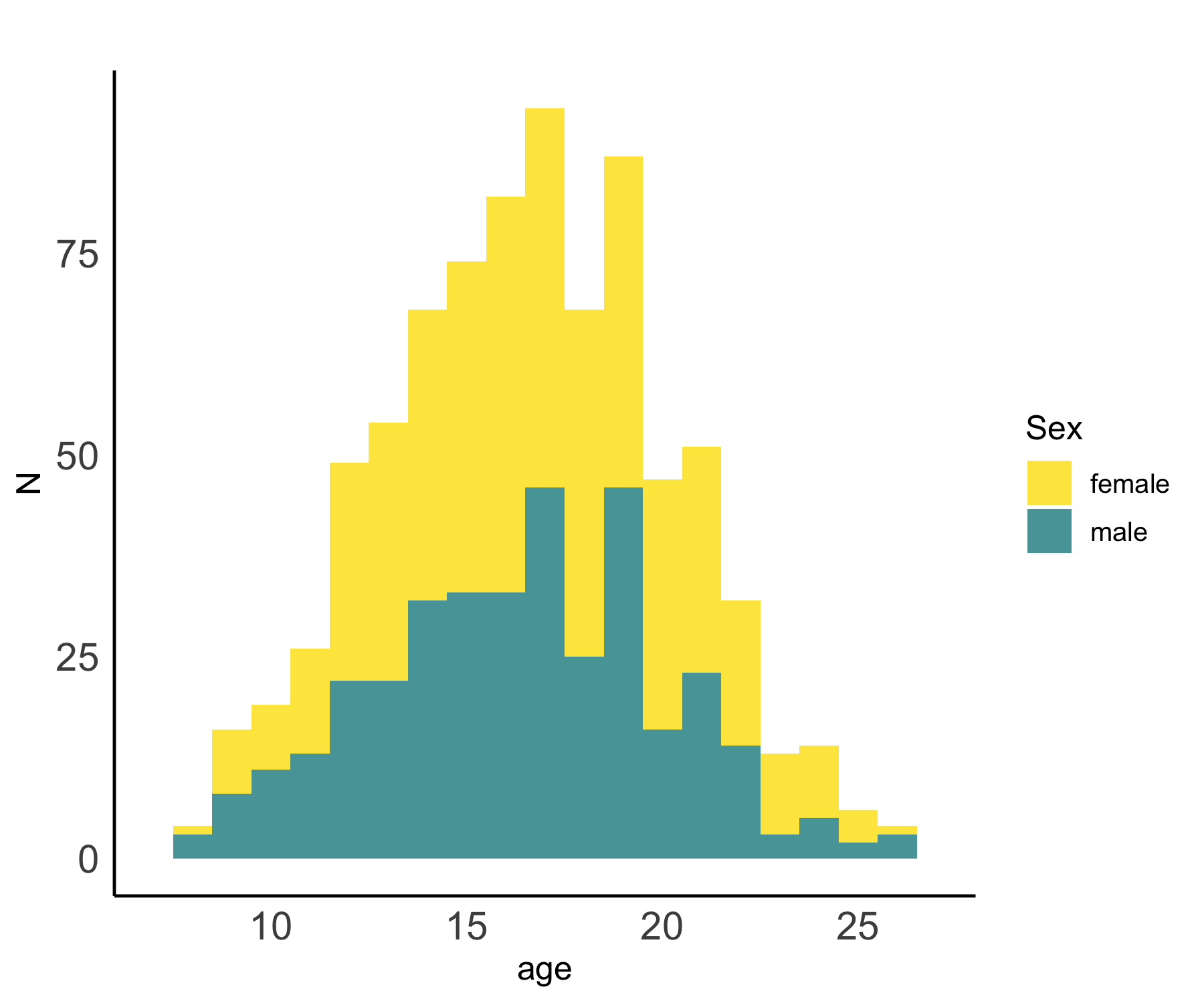

**SFigure 2.** Variability in annualized change over time for each individual (y-axis), against age at the midpoint of observation period. Purple line reflects group-level annualized change seen with age. The black dashed line marks 0 on the y-axis and the dashed gray lines represent the standard deviation of annualized change across the whole sample. A) Total gray matter volume, cortex volume, cortical thickness, white surface area, and white matter volume B) Subcortical gray matter volume and specific subcortical structures.

A)

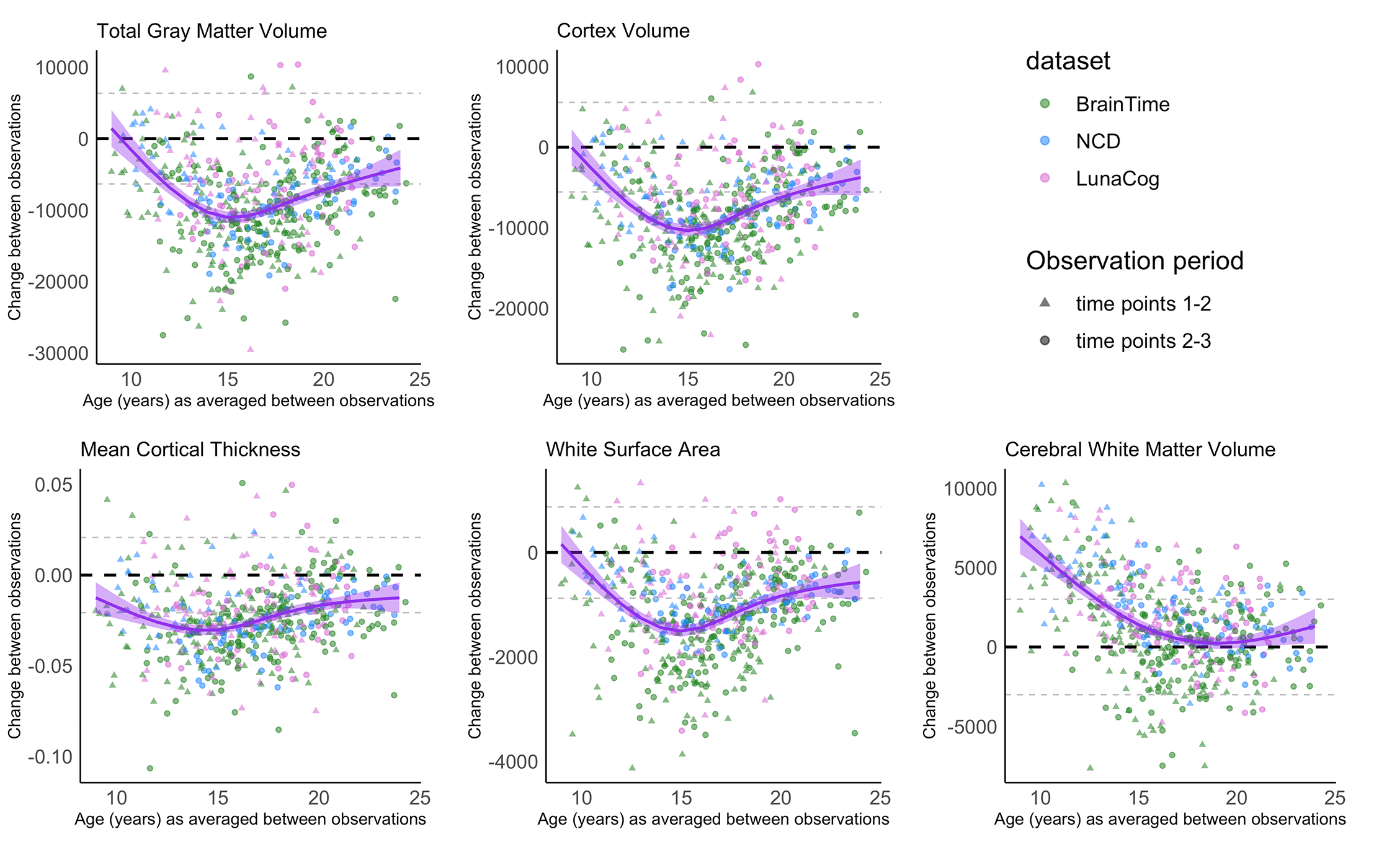

B)

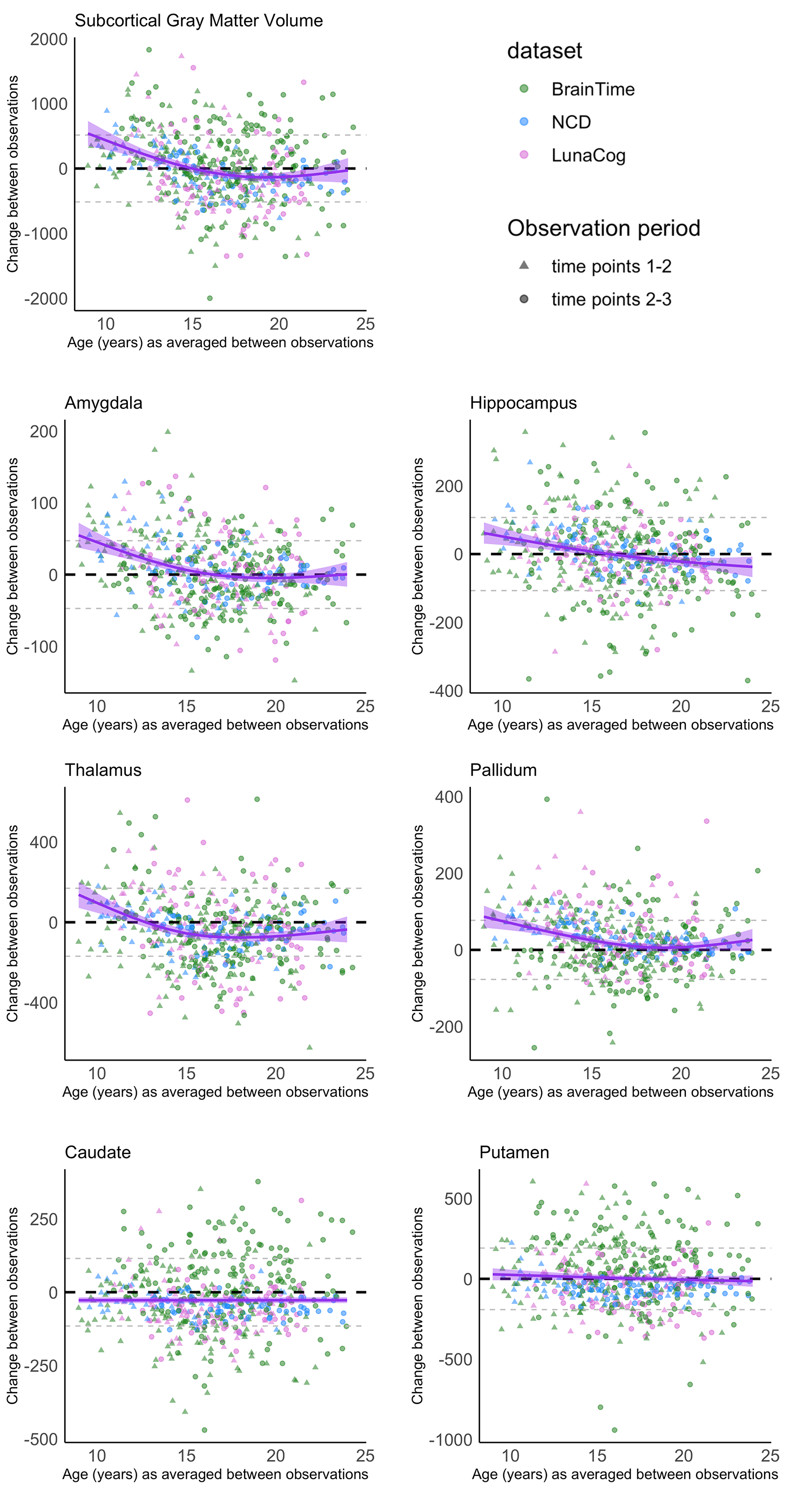

**SFigure 3.** Percent of individuals showing volume increases (purple), decreases (pink), or no change (turquoise) based on the individual’s age at the midpoint of observation period.

A)

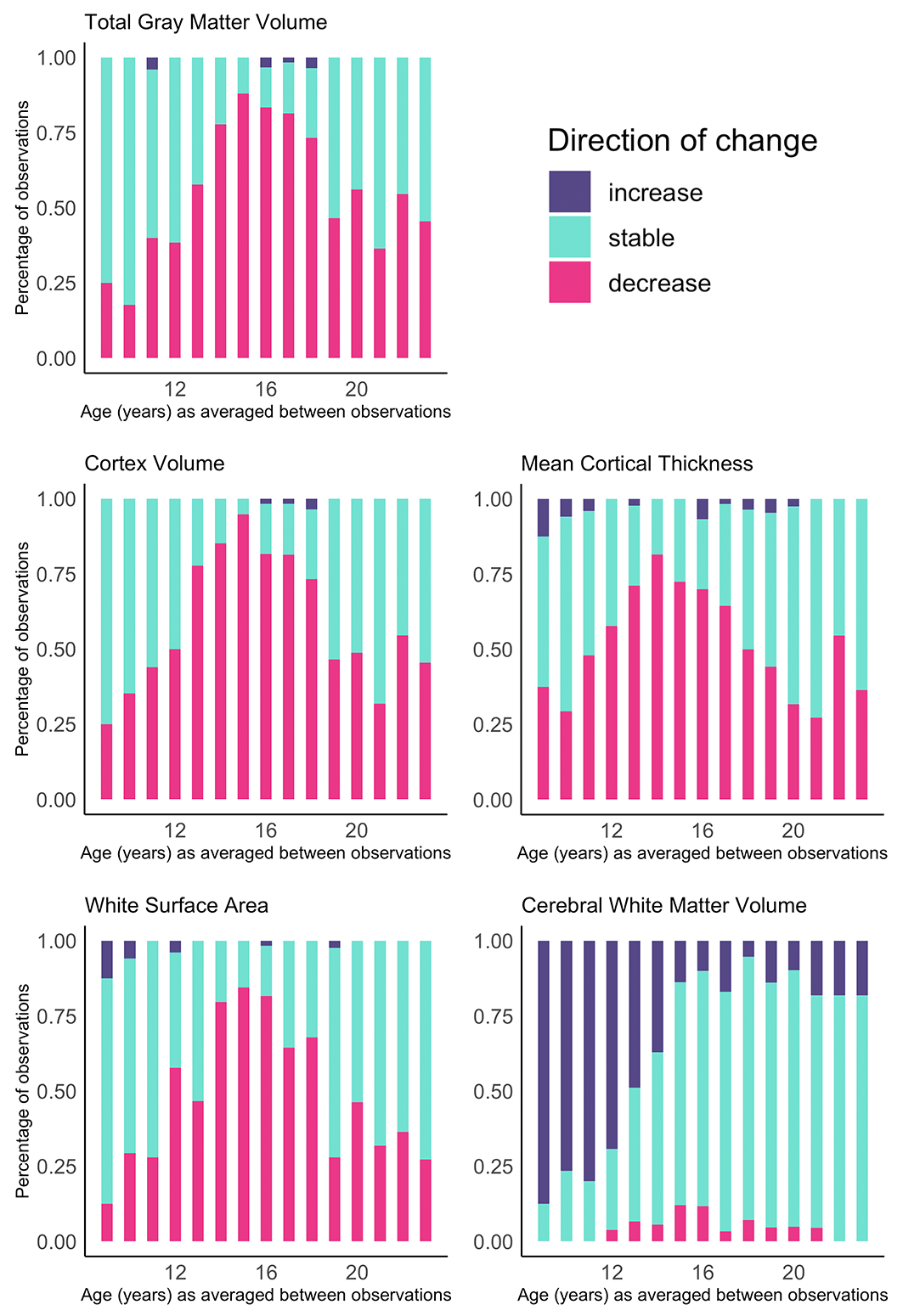

B)

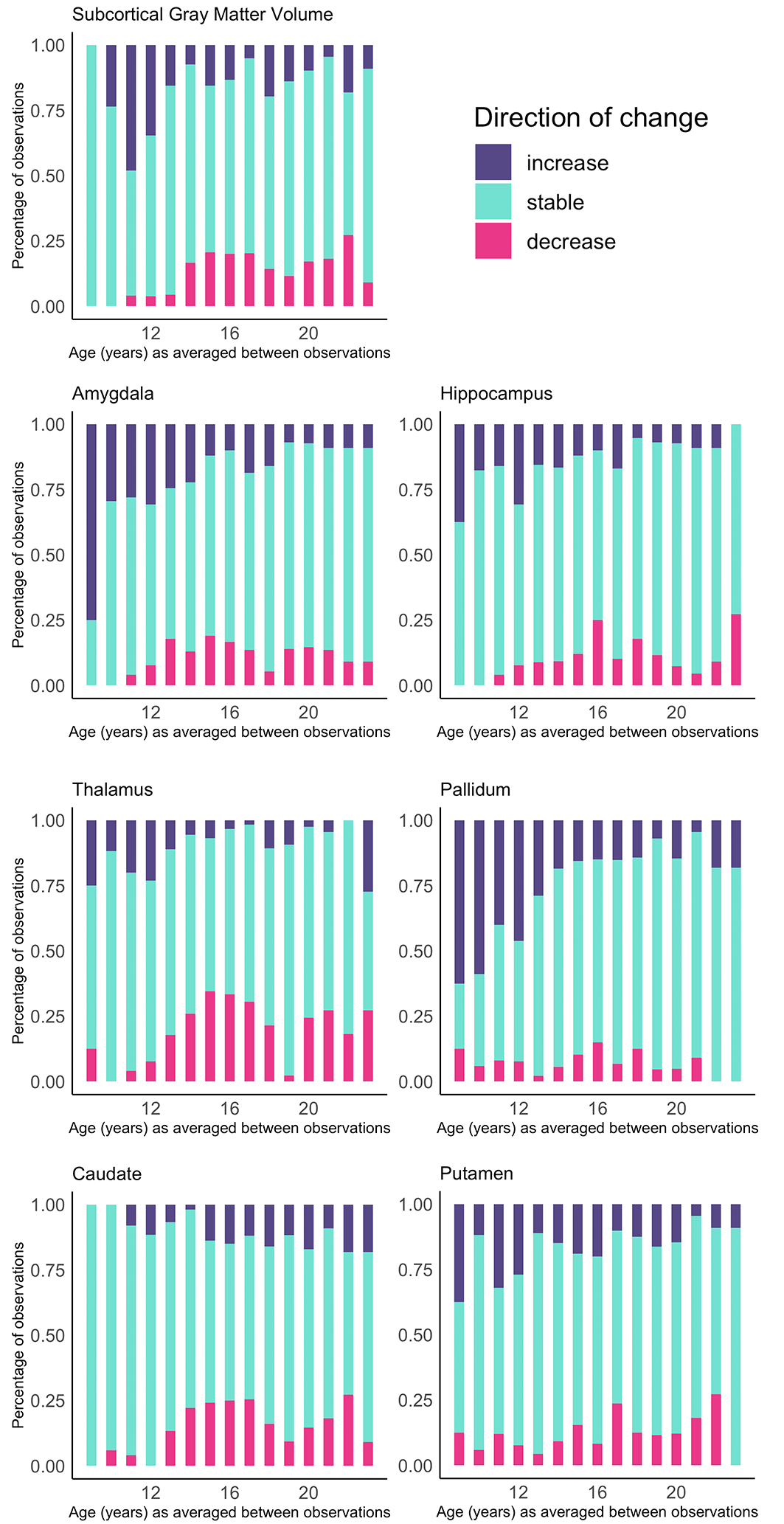
